# Genomic variations in SARS-CoV-2 genomes from Gujarat: Underlying role of variants in disease epidemiology

**DOI:** 10.1101/2020.07.10.197095

**Authors:** Madhvi Joshi, Apurvasinh Puvar, Dinesh Kumar, Afzal Ansari, Maharshi Pandya, Janvi Raval, Zarna Patel, Pinal Trivedi, Monika Gandhi, Labdhi Pandya, Komal Patel, Nitin Savaliya, Snehal Bagatharia, Sachin Kumar, Chaitanya Joshi

## Abstract

Humanity has seen numerous pandemics during its course of evolution. The list includes many such as measles, Ebola, SARS, MERS, etc. Latest edition to this pandemic list is COVID-19, caused by the novel coronavirus, SARS-CoV-2. As of 4th July 2020, COVID-19 has affected over 10 million people from 170+ countries, and 5,28,364 deaths. Genomic technologies have enabled us to understand the genomic constitution of the pathogens, their virulence, evolution, rate of mutations, etc. To date, more than 60,000 virus genomes have been deposited in the public depositories like GISAID and NCBI. While we are writing this, India is the 3rd most-affected country with COVID-19 with 0.6 million cases, and >18000 deaths. Gujarat is the fourth highest affected state with 5.44 percent death rate compared to national average of 2.8 percent.

Here, 361 SARS-CoV-2 genomes from across Gujarat have been sequenced and analyzed in order to understand its phylogenetic distribution and variants against global and national sequences. Further, variants were analyzed from diseased and recovered patients from Gujarat and the World to understand its role in pathogenesis. From missense mutations, found from Gujarat SARS-CoV-2 genomes, C28854T, deleterious mutation in nucleocapsid (N) gene was found to be significantly associated with mortality in patients. The other significant deleterious variant found in diseased patients from Gujarat and the world is G25563T, which is located in Orf3a and has a potential role in viral pathogenesis. SARS-CoV-2 genomes from Gujarat are forming distinct cluster under GH clade of GISAID.

## Introduction

As per the recent situation report-166 released by the World Health Organisation (WHO), as accessed on 4^th^ July 2020, total confirmed positive cases of COVID-19 across the globe are 10,922,324 resulting in 5,23,011 deaths. In many countries like China, Spain, Australia, Japan, South Korea, and USA, the second wave of SARS-CoV-2 infections has started **(Xu and Li 2020; Leung et al. 2020; Strzelecki 2020; Trade et al. 2020).** India is the third most affected country by COVID-19 after the USA and Brazil, with 6,48,315 cases and 18,655 deaths, respectively. Gujarat, located in the western part of India, is the fourth highest affected state in the world, with 36,123 cases and 1944 deaths. However, the death rate is 5.44%, which is almost two times higher than national average, with a recovery rate of 71.69% in the state of Gujarat, India. Therefore, understanding the pathogen evolution and virulence through genome sequencing will be key to understanding its diversity, variation and its effect on pathogenesis and disease severity. Global depositories like GISAID and NCBI databases are flooded with SARS-CoV-2 genomes with an average of 306 genomes per day being added from across the globe. SARS-CoV-2 genome size is 29 to 30.6 kb. The genome includes 10 genes which encode four structural and 16 non-structural proteins. Structural proteins are encoded by the four structural genes, including spike (S), envelope (E), membrane (M) and nucleocapsid (N) genes. The ORF1ab is the largest gene in SARS-CoV-2, which encodes the pp1ab protein and 15 non-structural proteins (nsps). The ORF1a gene encodes for pp1a protein, which also contains 10 nsps **(Shereen et al. 2020; Du et al. 2009)**.

In the present study, the whole genome of 361 SARS-CoV-2 from Gujarat have been sequenced and analyzed against 792 SARS-CoV-2 genomes across the globe, with the known patient status. The overall dataset comprises 277 confirmed positive COVID-19 patients, which included 100 females and 177 male patients. These genomes were studied against a total of 57,043 complete viral genome sequences as accessed on 4^th^ July 2020 to characterize their clades and variants distribution. Further statistical tools were applied to understand the differences in the variants with respect to disease epidemiology. In absence of the clinically approved drugs, vaccine, and possible therapy in treating COVID-19, tracking pathogen evolution through whole genome sequencing can be a very good tool in understanding the progression of pandemic locally as well as globally. This will also help in devising strategies for vaccine development, potential drug targets and host-pathogen interactions.

## Results

Samples were collected on the basis of COVID-19 incidence rate from across Gujarat from 13 different originating labs representing a total of 38 geographical locations from 18 districts of Gujarat, India **Supplemental Table 01**. The geographical distribution of the top three locations of viral isolates are represented by Ahmedabad (n=125), Surat (n=65), and Vadodara (n=53). Total 361 viral genomes from 277 patients have been sequenced in the study from which 132 were from females while 229 were from males. These patients were from 1 year to 86 years of age group with an average age of 47.80 yrs. Most of the COVID-19 positive patients had the symptoms of fever, diarrhoea, cough and breathing problems while some of them had the comorbid condition like hypertension and diabetes etc. The final outcome of these patients were classified as deceased, recovered, hospitalized or unknown status for further data analysis based on the available metadata information. These details are presented in **Supplemental Table S2.** Similarly, a data set of around 57,043 complete genomes of SARS-CoV-2 (up to 4^th^ July, 2020) downloaded from GISAID server and classified as per the patient status mentioned above. Chi-square test was performed to test the effect of gender and age group for Gujarat and global dataset. The female patients (at *p-value* 0.0240) in Gujarat dataset were observed to be at significantly higher death rate as compared to global dataset in deceased and recovered patients. The genomic dataset was further divided into different age groups of up to 40, 41-60 and over 60 years. The results indicated a significantly higher mortality rate at the age groups of 41-60 (at *p-value* 0.0391) and over 60 years in Gujarat (at *p-value* of 0.3932) compared against age groups in the global dataset. Mutation frequency profile of the Gujarat genome with the mutation spectrum is highlighted in **Figure 1**. including synonymous and missense mutations.

**Figure 1:**
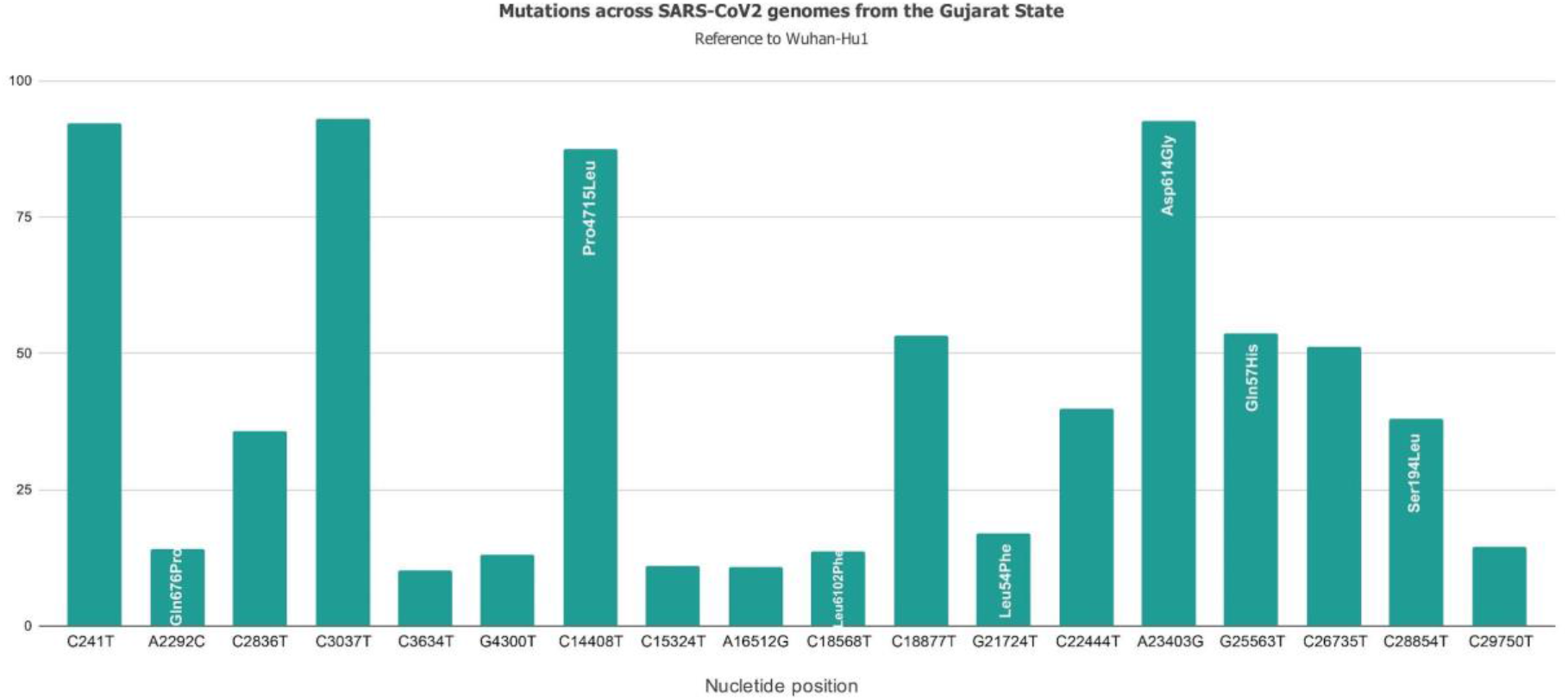
Mutation spectrum profile of 361 SARS-CoV-2 genome isolates sampled from 38 locations representing 18 districts of Gujarat, India including synonymous and missense mutation. The top mutations included C241T, C3037T, C14408T/Pro314Leu, C18877T, A23403G/Asp614Gly, G25563T/Gln57His and C26735T with frequency >50%.

### Genome sequencing

From a total of 277 patients, 84 had mixed infections. Mixed infections were judged by frequency of heterozygous mutations. Heterozygous mutation was considered only if it was supported by forward and reverse reads of an amplicon and 168 viral genomes were classified as two different haplotypes from 84 patients, and were observed with heterozygous allele frequencies and were manually divided in two genomes annotated with suffix “a” and “b”. All major alleles having read frequency ranging from 60 to 80 percent were included in “a” haplotypes while minor alleles having read frequency ranging from 20 to 40 percent were included in “b” haplotypes. Details of the reads, average coverage, mean read length, consensus genome length is provided in **Supplemental Table S2**.

### Phylogeny analysis

Phylogenetic analysis of 361 genomes were done as per the definitions of the PANGOLIN lineage and GISAID clades. The overall lineages distribution highlighted the dominant occurrence of B.1.36 (n=184), B.1 (n=143), A (n=14), B.6 (n=12), B.1.1 (n=5), B (n=3); while clade distribution highlights the dominant prevalence of GH (n=187), G (n=139), O (n=17), S (n=13), GR (n=4) and L (n=1) as mentioned in **Supplemental Table S3**. While none of the genomes from Gujarat belonged to clade V. In the global perspective, the distribution of the GISAID clades as on 4^th^ July 2020, from a total of 57,043 complete viral genome sequences, indicate the dominance of GR clade (n=15,784), G clade (n=12,541), GH clade (n=11,458), S clade (n=3,863), L clade (n=3,401), and V clade (n=3,640), where the “n” is the number of genomes. Maximum likelihood time-resolved phylogeny tree **Figure 2** using the TreeTime pipeline and Augur bioinformatics pipeline, annotated and visualized in the FigTree (**Rambaut et al., 2018; Hadfield et al. 2018**). Similarly, genomes classified into GISAID clades across the globe, and Gujarat are highlighted in **Figure 3**.

**Figure 2:**
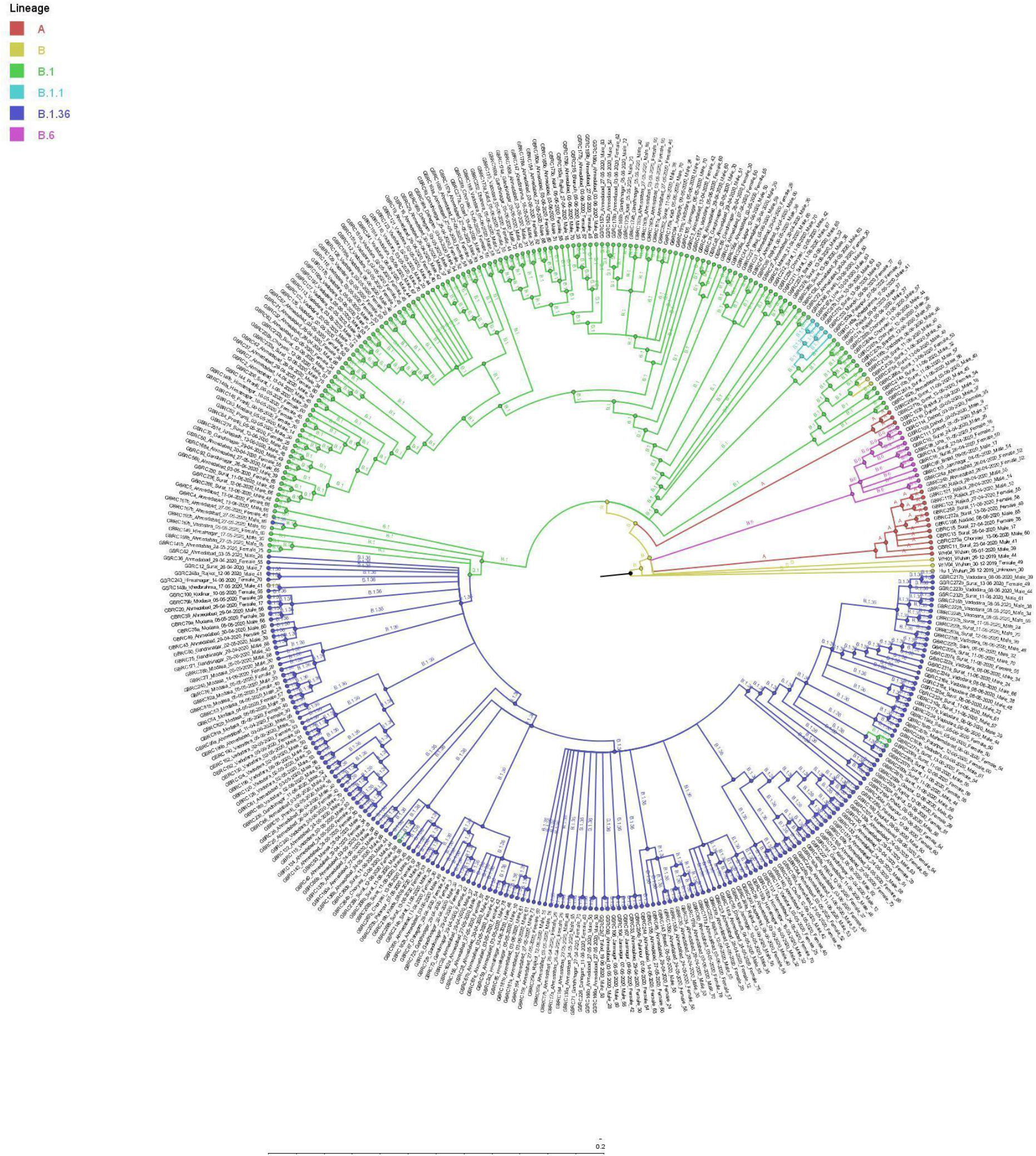
Phylogenetic distribution of lineage from 361 SARS-CoV-2 viral genomes of Gujarat, India with reference to the Wuhan/Hu-1/2019 (EPI_ISL_402125).

**Figure 3:**
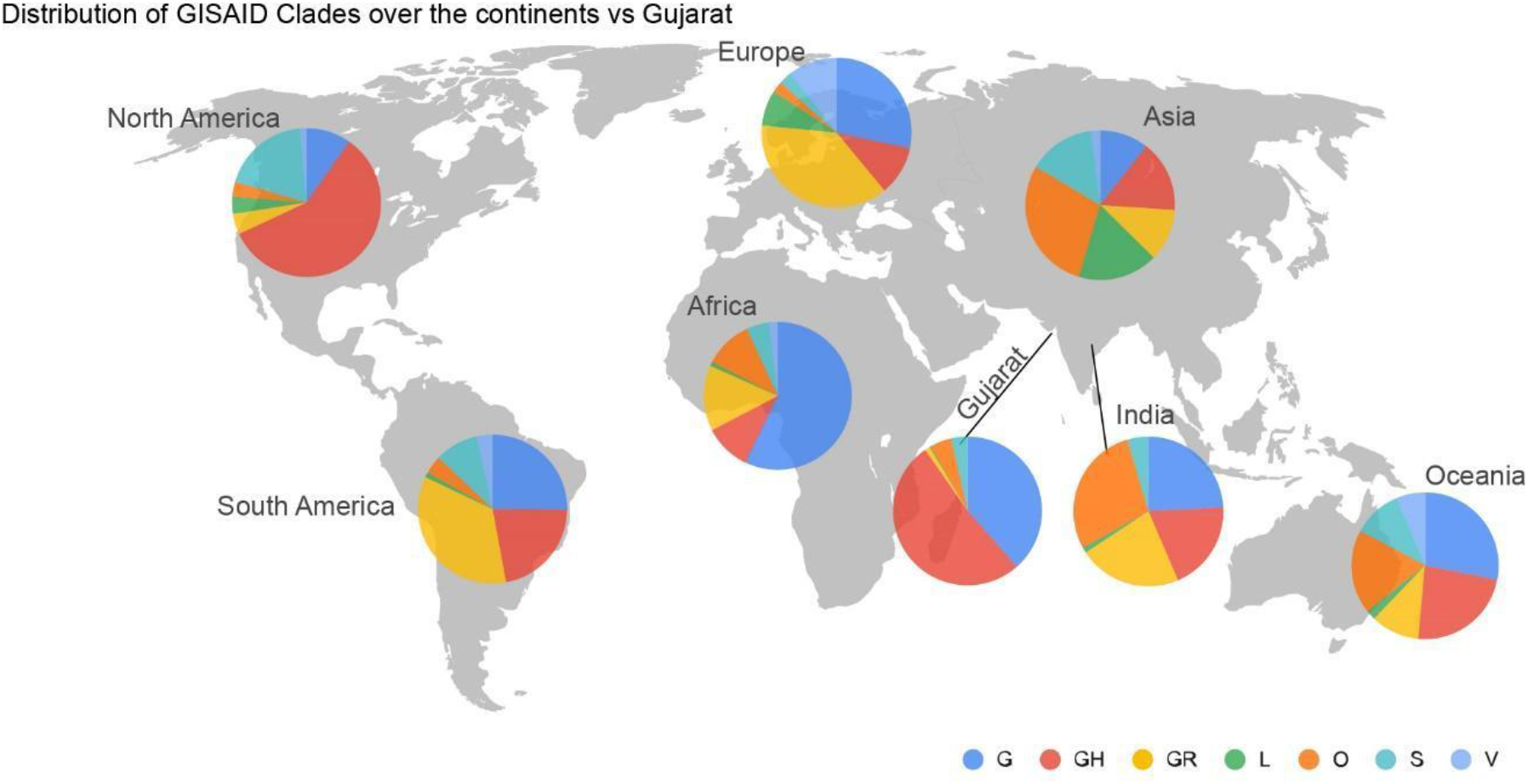
Distribution of the GISAID clades of the global genomes and Gujarat dataset as on 4th July 2020. Majority of the genomes from Gujarat cluster is dominated by prevalence of GH (n=187) and G (n=139) clades.

### Comparative analysis of mutation profile in SARS-CoV-2 genomes

To understand the significance of the mutations in the SARS-CoV-2 genome isolates from Gujarat, India we have analyzed and compared the mutation profile of the 361 viral isolates from Gujarat along with the global dataset obtained from GISAID with the known patient status of 753 viral genomes and 974 Indian genomes (unknown status). The bar chart displaying the comparative mutation analysis is represented as **Figure 4** displaying most frequently mutated regions on Gujarat SARS-CoV-2 genomes is described as in total, 23,711 mutations were observed in global sequences (n=57,043) of SARS-CoV-2 from GISAID where in 2,191 mutations were observed from 974 Indian isolates while 519 mutations were observed in genomes sequenced from Gujarat (n=361). Out of which 91 mutations were novel to Gujarat and 889 were novel to Indian genomes. A Venn diagram depicting mutations shared between sequences from Global, Indian and Gujarat isolates is given **Figure 5.** Similarly, comparison of the mutation profile analysis with *p-value* significance, frequency >5%, absolute count of the number of genomes with prevalence as represented in **Table 1**. Further frequencies of all the mutations were calculated by subtracting variants of Gujarat genomes from Indian and Global genomes with statistical significance.

**Table 1:** The overall comparison of missense and synonymous mutation frequency profile of Gujarat-361, India-974 and Global-57,043 dataset.

| NT position | AA position | Genome count |  |  | Frequency (%) |  |  | SIFT Score | Functional effect | p-value |
| --- | --- | --- | --- | --- | --- | --- | --- | --- | --- | --- |
|  |  | Gujarat<br>(n=361) | INDIA<br>(n=974) | Global<br>(n=57043) | Gujarat | INDIA | Global |  |  |  |
| C3037T |  | 336 | 512 | 42645 | 93.07 | 52.57 | 74.76 | 0.66 | Benign/Tolerated | 5.78682E-69 |
| A23403G | Asp614Gly | 334 | 520 | 42875 | 92.52 | 53.39 | 75.16 | 0.30 | Benign/Tolerated | 3.36085E-66 |
| C241T |  | 333 | 509 | 41904 | 92.24 | 52.26 | 73.46 | - | - | 5.89385E-63 |
| C14408T | Pro4715Leu | 316 | 500 | 42749 | 87.53 | 51.33 | 74.94 | 0.31 | Benign/Tolerated | 7.19968E-69 |
| G25563T | Gln57His | 194 | 75 | 12387 | 53.74 | 7.70 | 21.72 | 0.00 | Deleterious | 1.56659E-72 |
| C18877T |  | 192 | 66 | 1278 | 53.19 | 6.78 | 2.24 | 1.00 | Benign/Tolerated | 0 |
| C26735T |  | 185 | 67 | 361 | 51.25 | 6.88 | 0.63 | 1.00 | Benign/Tolerated | 0 |
| C22444T |  | 144 | 21 | 62 | 39.89 | 2.16 | 0.11 | 1.00 | Benign/Tolerated | 0 |
| C28854T | Ser194Leu | 137 | 24 | 937 | 37.95 | 2.46 | 1.64 | 0.05 | Deleterious | 0 |
| C2836T |  | 129 | 12 | 3 | 35.73 | 1.23 | 0.01 | 0.17 | Benign/Tolerated | 0 |
| G21724T | Leu54Phe | 61 | 1 | 198 | 16.90 | 0.10 | 0.35 | 0.69 | Benign/Tolerated | 0 |
| C29750T |  | 52 | 0 | 24 | 14.40 | 0.00 | 0.04 | #N/A | #N/A | 0 |
| A2292C | Gln676Pro | 51 | 0 | 0 | 14.13 | 0.00 | 0.00 | 0.05 | Deleterious | 0 |
| C18568T | Leu6102Phe | 49 | 0 | 38 | 13.57 | 0.00 | 0.07 | 0.01 | Deleterious | 0 |
| G4300T |  | 47 | 0 | 22 | 13.02 | 0.00 | 0.04 | 0.84 | Benign/Tolerated | 0 |
| C15324T |  | 40 | 33 | 1297 | 11.08 | 3.39 | 2.27 | 1.00 | Benign/Tolerated | 4.16974E-28 |
| A16512G |  | 39 | 0 | 8 | 10.80 | 0.00 | 0.01 | 1.00 | Benign/Tolerated | 0 |
| C3634T |  | 37 | 36 | 23 | 10.25 | 3.70 | 0.04 | 0.40 | Benign/Tolerated | 0 |
| G11083T | Leu3606Phe | 16 | 279 | 6059 | 4.43 | 28.64 | 10.62 | 0.01 | Deleterious | 9.42938E-74 |
| C28311T | Pro13Leu | 13 | 255 | 689 | 3.60 | 26.18 | 1.21 | 0.00 | Deleterious | 0 |
| C6312A | Thr2016Lys | 12 | 259 | 531 | 3.32 | 26.59 | 0.93 | 0.03 | Deleterious | 0 |
| C23929T |  | 12 | 238 | 493 | 3.32 | 24.44 | 0.86 | 1.00 | Benign/Tolerated | 0 |
| C13730T | Ala4489Val | 11 | 277 | 660 | 3.05 | 28.44 | 1.16 | 0.00 | Deleterious | 0 |
| C5700A | Ala1812Asp | 7 | 140 | 0 | 1.94 | 14.37 | 0.00 | 0.38 | Benign/Tolerated | 0 |
| C313T |  | 4 | 146 | 880 | 1.11 | 14.99 | 1.54 | 0.84 | Benign/Tolerated | 7.1703E-218 |
| C1059T | Thr265Ile | 2 | 7 | 9719 | 0.55 | 0.72 | 17.04 | 0.03 | Deleterious | 2.39204E-55 |

**Figure 4:**
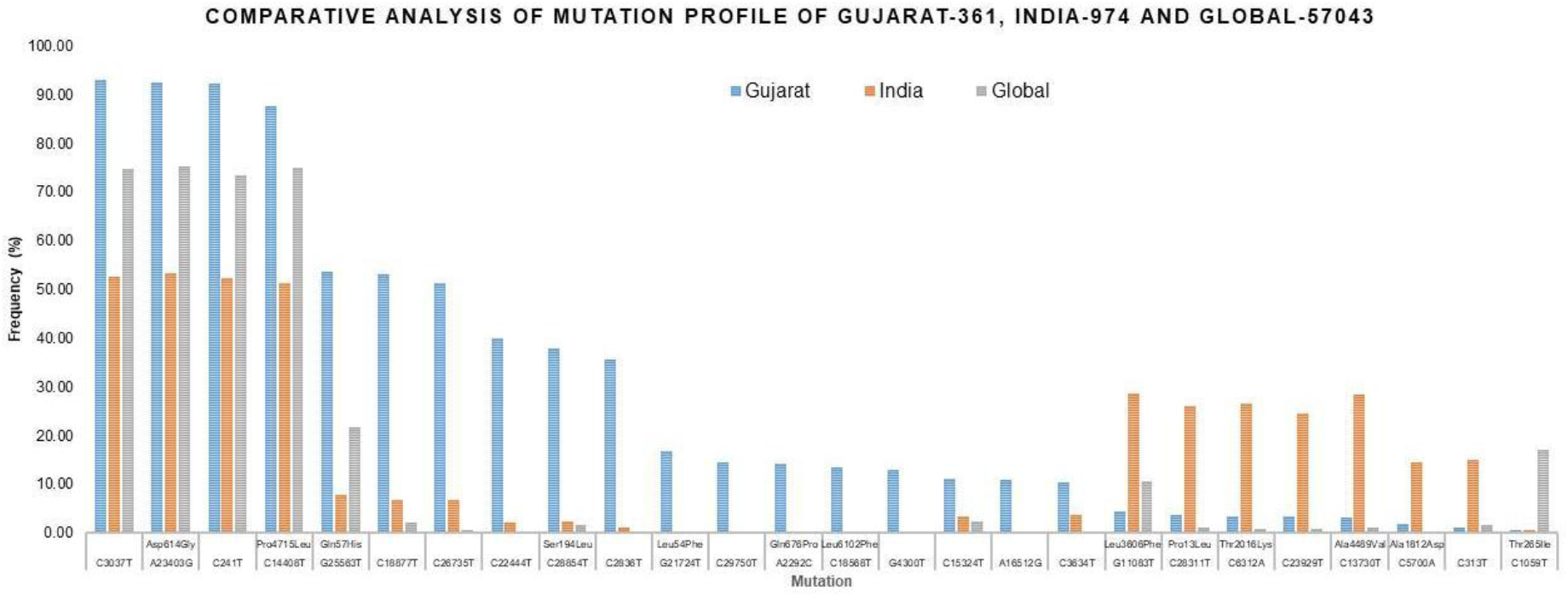
Synonymous and missense mutation profile of the Gujarat-361, India-974, and Global-57043 dataset with >5% frequency.

**Figure 5:**
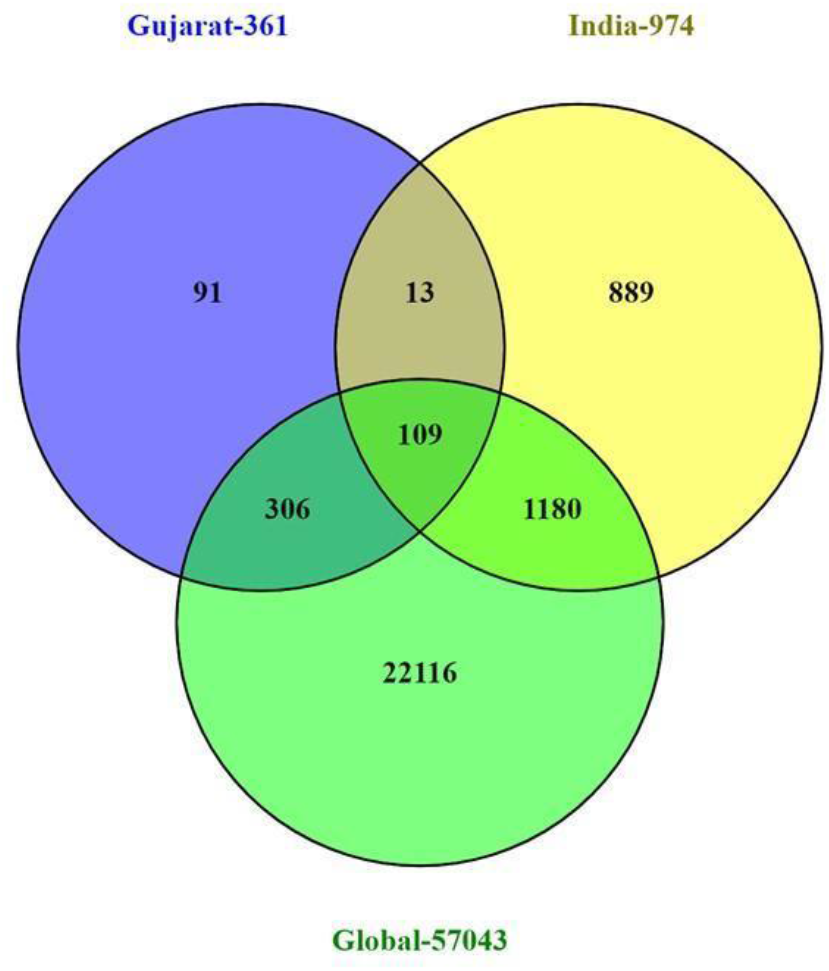
Venn diagram representing the mutually common and exclusive synonymous and missense mutations among the Gujarat-361, India-974, and Global-57,043 dataset.

Mutations C241T, C3037T, A23403G and C14408T mutations were at higher frequencies (>50%) in all the genomes while G11083T, C13730T, C28311T, C6312A and C23929T mutations were predominated (>24% frequency) in Indian genomes however at very low frequency (<15%) in comparison with Global and Gujarat genomes. The mutations G25563T, C26735T, C18877T at frequency >51% while C2836T, C22444T and C28854T at >35% frequency and G21724T, C29750T, C18568T and A2292C were occurring at >13% frequency from genomes sequences of Gujarat. All these mutations were found to be statistically significant at *p-value* <0.001. Out of these mutations A23403G, C14408T, G25563T and C28854T were missense mutations. The detailed mutation frequency profile is provided as **Supplemental Table S4**. With reference to Indian genomes, G11083T, C28311T, C6312A, C23929T and C13730T were found to be occurring at more than 24% frequencies (*p-value* <0.001). From these mutations, G11083T, C28311T and C6312A were found to be missense mutations. G11083T and C6312A lie in the region of Orf1a encoding Nsp6. Further deceased versus recovered patient mutation profile analysis of the known patient’s status dataset from Gujarat and Global is represented in **Figure 6**. Similarly, comparison of missense mutation profile of deceased verses recovered patients with genome count, frequency >5%, and *p-value* for global dataset is represented in **Table 2** and for Gujarat dataset **Table 3**.

**Table 2:** Comparison of missense mutation frequency in deceased vs recovered patients from global dataset.

| NT mutation | AA mutation | Global mutation count (genomes) |  | Global frequency (%) |  | SIFT Score | Functional effect | p-value |
| --- | --- | --- | --- | --- | --- | --- | --- | --- |
|  |  | Deceased (n=131) | Recovered (n=622) | Deceased | Recovered |  |  |  |
| A23403G | Asp614Gly | 117 | 378 | 89.31 | 60.77 | 0.3 | Benign/Tolerated | 3.95251E-10 |
| C14408T | Pro4715Leu | 115 | 374 | 87.79 | 60.13 | 0.31 | Benign/Tolerated | 1.64386E-09 |
| G25563T | Gln57His | 32 | 131 | 24.43 | 21.06 | 0 | Deleterious | 0.395157807 |
| C1059T | Thr265Ile | 18 | 95 | 13.74 | 15.27 | 0.03 | Deleterious | 0.655252139 |
| G25088T | Val1176Phe | 27 | 0 | 20.61 | 0.00 | #N/A | #N/A | 9.19649E-31 |
| C12053T | Leu3930Phe | 11 | 0 | 8.40 | 0.00 | 0.00 | Deleterious | 3.3299E-13 |
| G11083T | Leu3606Phe | 6 | 48 | 4.58 | 7.72 | 0.01 | Deleterious | 0.205974179 |
| T28144C | Leu84Ser | 3 | 92 | 2.29 | 14.79 | 0.37 | Benign/Tolerated | 8.98524E-05 |
| G26144T | Gly251Val | 1 | 39 | 0.76 | 6.27 | 0 | Deleterious | 0.010644459 |
| C28833T | Ser187Leu | 0 | 31 | 0.00 | 4.98 | 0.00 | Deleterious | 0.009068597 |

**Table 3:** Comparison of missense mutation frequency in deceased vs recovered patients from Gujarat dataset.

| NT mutation | AA mutation | Gujarat mutation count (genomes) |  | Frequency (%) |  | SIFT Score | Functional effect | p-value |
| --- | --- | --- | --- | --- | --- | --- | --- | --- |
|  |  | Deceased (n=43) | Recovered (n=172) | Deceased | Recovered |  |  |  |
| A23403G | Asp614Gly | 42 | 160 | 97.67 | 93.02 | 0.30 | Benign/Tolerated | 0.252398792 |
| C14408T | Pro4715Leu | 41 | 154 | 95.35 | 89.53 | 0.31 | Benign/Tolerated | 0.240407116 |
| G25563T | Gln57His | 23 | 94 | 53.49 | 54.65 | 0.00 | Deleterious | 0.891082474 |
| C28854T | Ser194Leu | 19 | 59 | 44.19 | 34.30 | 0.00 | Deleterious | 0.227942491 |
| G16078A | Val5272Ile | 4 | 9 | 9.30 | 5.23 | 0.00 | Deleterious | 0.316597442 |
| C23277T | Thr572Ile | 4 | 5 | 9.30 | 2.91 | 0.57 | Benign/Tolerated | 0.061074771 |
| G21724T | Leu54Phe | 1 | 28 | 2.33 | 16.28 | 0.69 | Benign/Tolerated | 0.016585523 |
| A2292C | Gln676Pro | 1 | 24 | 2.33 | 13.95 | 0.05 | Deleterious | 0.0333774 |
| C18568T | Leu6102Phe | 1 | 24 | 2.33 | 13.95 | 0.01 | Deleterious | 0.0333774 |
| G28878A | Ser202Asn | 0 | 12 | 0.00 | 6.98 | 0.00 | Deleterious | 0.074666193 |

**Figure 6:**
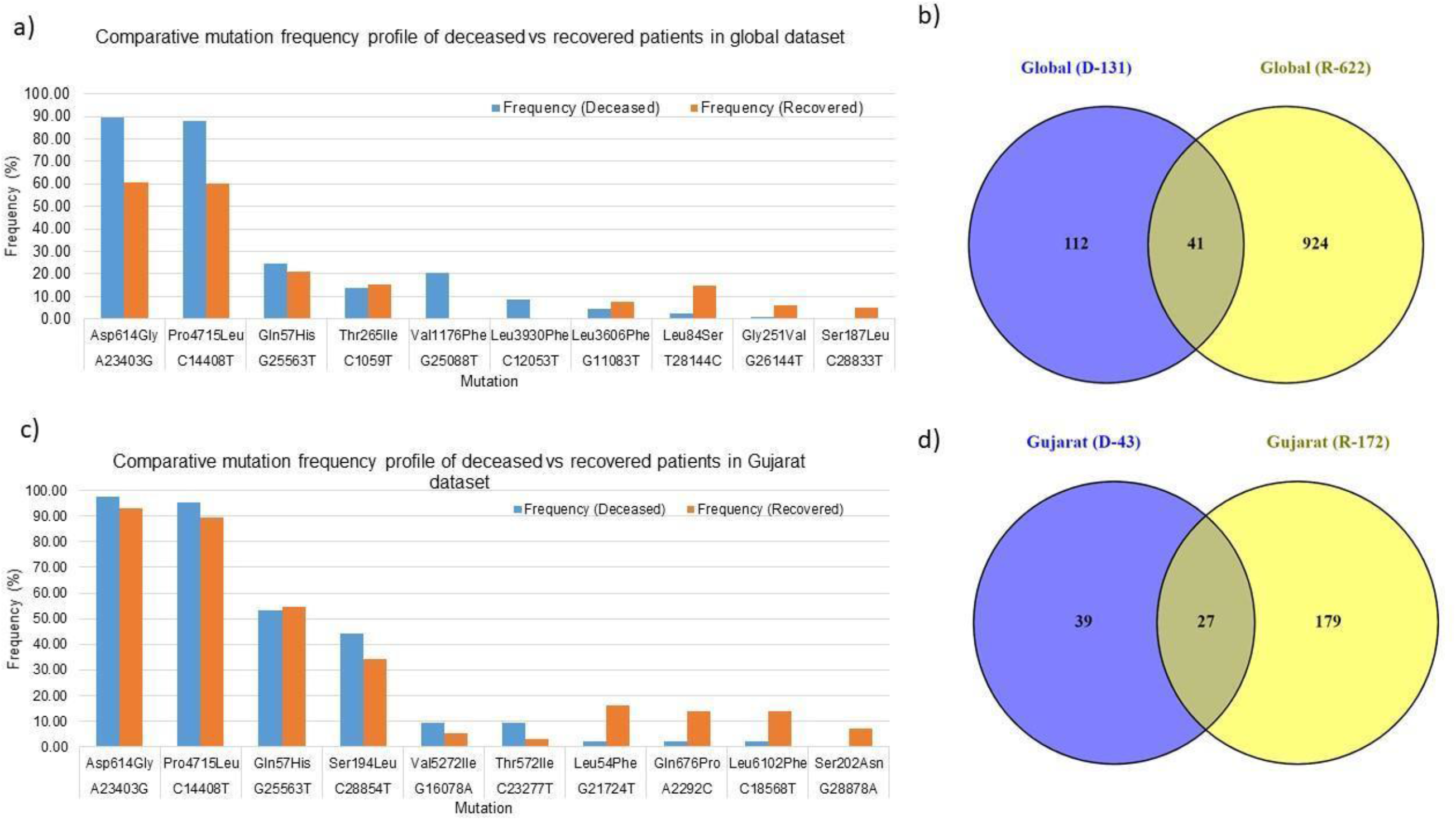
Global and Gujarat mutation frequency analysis of missense mutations **a)** Bar chart for global deceased versus recovered patients **b)** Venn diagram of the global deceased versus recovered patients **c)** Bar chart for global deceased versus recovered patients **d)** Venn diagram of the Gujarat deceased versus recovered patients. Additional Supplemental Table S5 and S6 provided for details of the missense mutations in Gujarat and Global dataset of deceased versus recovered patients.

The statistical significance association of the gender and age of the deceased and recovered patients from Gujarat and global dataset revealed the significant *p-value* for female patients in both datasets considered for analysis. Similarly, for age group 41-60 yrs. highlighted the higher observation of death rate in patients with known status as given in **Table 4**.

**Table 4:** Chi-square test analysis of the deceased and recovered patients for gender and age group.

|  |  | Gujarat (n=361) |  | Global (n=57043) |  |  |
| --- | --- | --- | --- | --- | --- | --- |
|  |  | Deceased | Recovered | Deceased | Recovered | *p-value* |
| **Total Sample** |  | **43** | **172** | **131** | **622** | **0.380671969** |
| **Gender** | **Male** | **24** | **113** | **86** | **383** | **0.826887912** |
|  | **Female** | **19** | **59** | **45** | **278** | **0.024025225** |
| **Age (Yrs)** | **0-40** | **2** | **70** | **8** | **288** | **0.971968847** |
|  | **41-60** | **19** | **77** | **30** | **234** | **0.039186032** |
|  | **>60** | **22** | **25** | **93** | **139** | **0.393230893** |

## Discussion

SARS-CoV-2 viral genome analysis from Gujarat highlights the distinct genomic attributes, geographical distribution, age composition and gender classification. These features also highlight unique genomic patterns in terms of synonymous and non-synonymous variants associated with the prevalence of dominant clades and lineages with distinct geographical locations in Gujarat. This work also highlights the most comprehensive genomic resources available so far from India. Identifying variants specific to the deceased and recovered patients would certainly aid in better treatment and COVID-19 containment strategy. The fatality rate compared with different geographical locations may point towards the higher virulence profile of certain viral strains with lethal genetic mutations, but this remains clinically unestablished. Perhaps the onset of clinical features in the symptomatic patients help in prioritizing the diagnosis and testing strategy.

Genomes reported from India are having diverse mutation profiles. The first case report of complete genome sequence information from India is from a patient in Kerala with a direct travel history to Wuhan, China. Similarly, other isolates from India cluster with Iran, Italy, Spain, England, USA and Belgium and probably similar isolates are transmitting in India and may also have variable mutation profile **(Mondal et al.; Yadav et al. 2020; Potdar et al. 2020).** The dominance of a particular lineage or clade at a particular location merely does not establish the biological function of the virus type isolate in terms of higher death rate but the epidemiological factors such as clinically diagnosed co-morbidity, age, gender or asymptomatic transmission most likely influencing factor in transmission. Sampling biases could certainly influence the prediction models but it would definitely narrow down to particular types of isolates and unique mutations which further experimentally validated to establish their clinical significance. Further, in subsequent analysis, we have also analysed and identified the mortality rate in different age groups revealing the age group of 41-60 years were statistically significant with *p-value* of 0.03901.

The geographic distribution of the viral isolates is denoted in the phylogeny with the maximum SARS-CoV-2 positive samples sequenced from Ahmedabad (n=125), followed by Surat (n=65), Vadodara (n=53), Gandhinagar (n=28), Sabarkantha (n=18) and Rajkot (n=18). The distribution of dominant lineages in Ahmedabad is steered by occurrences of B.1.36 (n=72), B.1 (n=51) and B.6 (n=2). The concept of lineages, clades, haplotypes or genotypes is slightly perplexing and overlapping in terms of definitions with respect to different depositories and analytics. Therefore, it is most important to define mutations in the isolates that determine their unique position in phylogeny with respect to geographical distribution, age, gender, and locations of the genotypes etc. Phylogenetic distribution of the viral genomes across different geographical locations along with metadata information should help in evaluation of the posterior distribution, virulence, divergence times and evolutionary rates in viral populations **(Drummond and Rambaut 2007).** The recurrent mutations occurring independently multiple times in the viral genomes are hallmarks of convergent evolution in viral genomes with significance in host adaptability, spread and transmission. Even though, contested in terms of mechanisms driving the pathogenicity and virulence across different hosts and specifically to human populations across different geographical locations **(van Dorp et al. 2020; Grifoni et al. 2020)**.

### Incidence of mutations in deceased and recovered patients

In the context of the globally prevalent mutations across different geographical locations, we have analysed viral genome isolates with most frequent mutations present in the patients from those who have suffered casualties. The higher death rate, especially in Ahmedabad, India became a cause of serious concern and remains elusive to be identified with enough scientific evidence. We have identified differentially dominant and statistically significant mutations prevalent in the viral genome isolates in Gujarat, India. The dominant mutations in the deceased patients were represented by the change in A23403G was observed at a frequency of 97.67% in Gujarat (*p-value* of 0.2523) and 89.31% frequency in global genomes with known patient status (*p-value* of <0.00001). These missense mutations are found to be observed in the spike protein of the coronavirus genome. The well-known function of the viral spike protein is in mediating the infection by interacting with the Angiotensin-converting enzyme 2 (ACE2) receptor **(Guo et al. 2020; Li et al. 2005; Chu et al. 2020; Guan et al. 2020)** of the human host species. Another mutation, C14408T with a frequency of 95.35%, present in the Orf1b gene encoding RNA directed RNA polymerase (RDRP) non-structural protein (nsp12) with a *p-value* of 0.2404 in deceased versus recovered patients from Gujarat, while also being observed statistically significant in the global dataset with a *p-value* of <0.00001 with a frequency of 87.79 percent. The comparative analysis of the patients deceased (n=43) and recovered patients (n=172) in Gujarat as highlighted in **Figure 6** as represented in Venn diagram. In contrast, the functional role of the RDRP enzyme activity is necessary for the viral genome replication and transcription of most RNA viruses (**Imbert et al. 2006; Velazquez-Salinas et al., 2020**).

The exclusive dominant mutations present in the population of Gujarat, India, and those simultaneously being statistically significant were at G25563T and at C28854T were present in the Orf3a and N gene, respectively. The Orf3a gene encodes a protein involved in the regulation of inflammation, antiviral responses, and apoptosis. Mutation in these regions alters the functional profile of the nuclear factor-kβ (NF-kβ) activation and (nucleotide-binding domain leucine-rich repeat-containing) NLRP3 inflammasome. One of the main features of Orf3a protein is having the presence of a cysteine-rich domain, which participates in the enzymatic nucleophilic substitution reactions. This protein is expressed abundantly in infected and transfected cells, which localizes to intracellular and plasma membranes and also induces apoptosis in transfected and infected cells **(Issa et al. 2020).** This enzyme mediates extensive proteolytic processing of two overlapping replicase polyproteins, pp1a and pp1ab, to yield the corresponding functional polypeptides that are essential for coronavirus replication and transcription processes **(Kohlmeier and Woodland 2009; Benvenuto et al. 2020)**. Whereas, in case of mutation at position C28311T leading to change of amino acid proline to leucine lies in the nucleocapsid (N) gene which has a role in virion assembly and release and plays a significant role in the formation of replication-transcription complexes **(Yin 2020; J Alsaadi and Jones 2019; Liu 2019; Wu et al. 2020)**. Similarly, the nucleocapsid (N) protein is a highly basic protein that could modulate viral RNA synthesis **(Millet and Whittaker 2015; Hassan et al. 2020; Sarif Hassan et al. 2020)**. The Sorting Intolerant from Tolerant (SIFT) scores of these mutations were determined and also signifies the functional effect change in whether an amino acid substitution affects protein function or not in terms of the deleterious effect or benign tolerated. The SIFT score ranges between 0.0 to 0.05 (deleterious) and 0.05 to 1.0 (tolerated) to differentiate the mutation effect **(Vaser et al. 2016)**. The predicted SIFT score of the mutation G25563T in the Orf3a and C28854T in the N gene was classified to be deleterious in nature. Similarly, a comparison analysis of the global (n=57,043), India (n=974) and Gujarat (n=361), where the “n” is the number of genomes included in the analysis indicates the overall dominance of C241T, C3037T, A23403G, C14408T, and G25563T. Furthermore, suggestive of the comparative dominant mutation profile, including nonsynonymous and missense mutations.

Analysis of the dataset of the global deceased (n=131) and recovered patients (n=622) with known status from the metadata information available on GISAID server with the complete genome sequences considered in the analysis indicates the dominance of the A23403G, C14408T and G25563T. The overall comparison of the mutation profile of the patient dataset of deceased and recovered samples is highlighted in **Figure 8**. While comparing the exclusive missense mutation profile of the patients recovered and deceased in Gujarat (x=43, y=172) and global dataset (x=131, y=622), where the “x” is number of deceased patients and “y” is number of recovered patients.

While analyzing missense variants from global and Gujarat dataset among deceased and recovered patients, identified four major mutations to be significantly associated with deceased patients. However, in the context of global dataset mutation C14408T in the RdRp gene and A23403G in spike protein gene were found to be associated with deceased patients at *p-value* <0.00001. Mutations in the N gene at C28854T and mutation in Orf3a at G25563T gene from diseased patients in Gujarat were found to be significant among deceased patients at *p-value* 0.0094 and 0.231 respectively. Moreover, C28854T is forming a distinct sub-cluster under 20A (A2a as per old classification of next strain) clade with a frequency of 37.95, 2.46 and 1.64 percent (*p-value* <0.01) in Gujarat, India and global dataset respectively. The same is highlighted in **Figure 7**. The same is proposed as a new sub-clade 20D in the next strain and GHJ in GISAID. This sub-clade is also present in genomes sequenced from Bangladesh and Saudi Arabia. Both these proteins have a significant role in viral replication and pathogenesis **(Pachetti et al. 2020; Luan et al. 2020; Peter and Schug)**.

**Figure 7:**
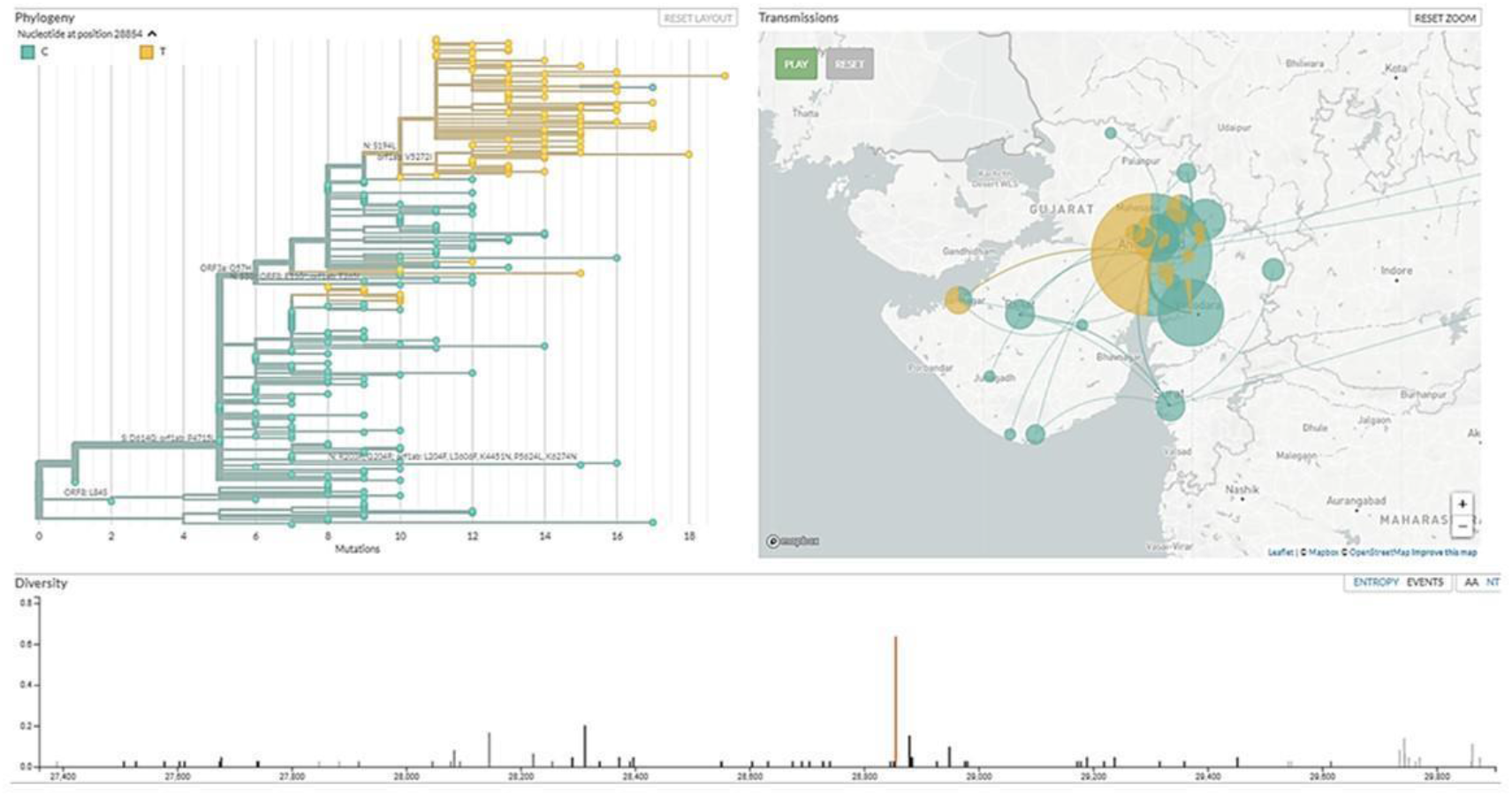
Distinct cluster of the C28854T genomes in Gujarat SARS-CoV-2 viral genome isolates.

**Figure 8:**
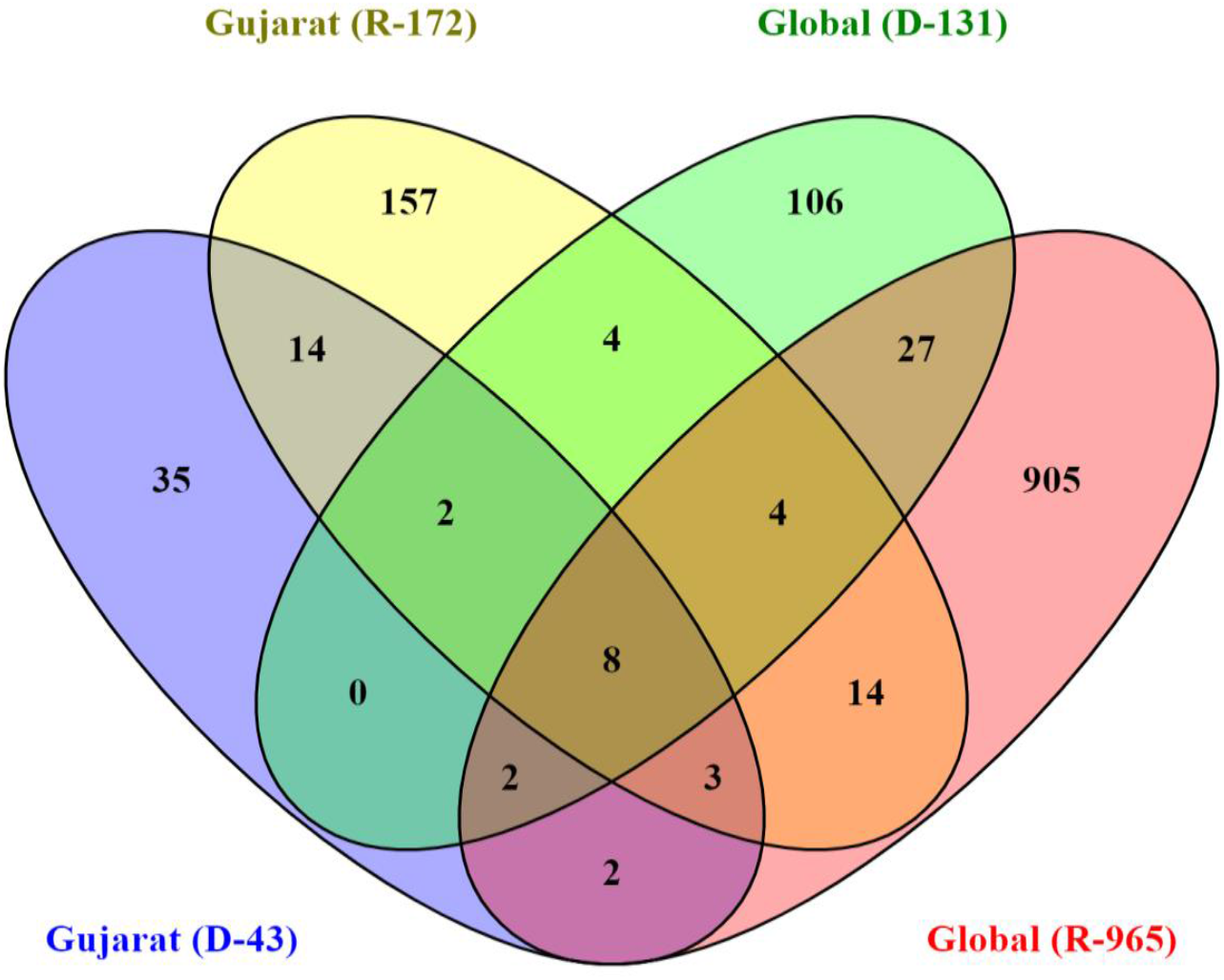
Overall comparison of the missense mutations in Gujarat (R=172, D=43), Global (R=622, D=131) where “R” is number of genomes from recovered patients, and “D” is the number of genomes from deceased patients.

The association of the mutations with the viral transmission and mortality rate remains a mystifying puzzle for the global scientific community. The identification and validation of these mutations should pave the way forward for the development of treatment and diagnostics of coronavirus disease. The evading host immune response and defence mechanism sufficiently improve the adaptive behaviour of pathogenic species, thus, making them highly contagious. Further, laboratory and experimental studies need to be carried out to validate the exact role of this particular mutation in respect to the molecular pathways and interactions in the biological systems despite being a strong possible mutation candidate found in the Gujarat region.

## Conclusion

The genomics approach has been a useful resource to identify and characterize the virulence, pathogenicity, and host adaptability of the sequenced viral genomes. Identification and characterization of the frequently mutated positions in SARS-CoV-2 genome will certainly help in the further understanding of infection biology of the coronaviruses, development of vaccines and therapeutics, and drugs repurpose candidates using predictive computational biology resources. The present study highlights the genomic signature and mutation profile of the 361 SARS-CoV-2 viral genome isolates from 38 different locations representing 18 districts across Gujarat, India. Further, we have reported significant variants associated with mortality in Gujarat and Global genomes. As the pandemic is progressing, the virus is also diverging into different clades. This also provides adaptive advantages to viruses in progression of the disease and its pandemic potential. In this study, we have reported a distinct cluster of coronavirus under 20A clade of Nextstrain and proposed it as 20D as per next strain analysis or GHJ as per GISAID analysis, predominantly present in Gujarat genomes. Understanding the pathogen genome and tracking its evolution will help in devising better strategies for the development of diagnosis, treatment, vaccine in response to pandemic.

## Material and methods

### Sample collection and processing

Nasopharyngeal and oropharyngeal swabs from a total of 277 individuals tested positive for COVID-19 from 38 locations representing 18 districts of Gujarat were collected after obtaining informed consent and appropriate ethics approval. The numbers of samples from these locations were selected on the basis of disease spread in Gujarat. The details of the samples collected from each location is shown in **Supplemental Table S1**. Samples were transported as per standard operating procedures as prescribed by the World Health Organisation (WHO) and Indian Council of Medical Research (ICMR, New Delhi; SoP No: ICMR-NIV/2019-nCoV/Specimens_01) to research lab at GBRC and further stored at −20° C till processed.

### Whole genome sequencing of SARS-CoV-2

Total genomic RNA from the samples were isolated using QIAamp Viral RNA Mini Kit (Cat. No. 52904, Qiagen, Germany) following prescribed biosafety procedure. cDNA from the extracted RNA was made using SuperScript™ III Reverse Transcriptase first strand kit (Cat. No: 18080093, ThermoFisher Scientific, USA) as per the procedures prescribed. SARS-CoV-2 genome was amplified by using Ion AmpliSeq SARS-CoV-2 Research Panel (ThermoFisher Scientific, USA) that consists of two pools with amplicons ranging from 125 bp to 275 bp in length and covering >99% of the SARS-CoV-2 genome, including all serotypes. Amplicon libraries were prepared using Ion AmpliSeq™ Library Kit Plus (Cat. No: A35907; ThermoFisher Scientific, USA). These libraries were quantified using the Ion Library TaqMan™ Quantitation Kit (Cat. No: 4468802, ThermoFisher Scientific, USA). The quality of the library was checked on DNA high sensitivity assay kit on Bio-analyser 2100 (Agilent Technologies, USA) and were sequenced on the Ion S5 Plus sequencing platform using 530 chip.

### Raw data quality assessment and filtering

Quality of data was assessed using FASTQC v. 0.11.5 (**Andrews et al., 2014**) toolkit. All raw data sequences were processed using PRINSEQ-lite v.0.20.4 (**Schmieder and Edwards 2011**) program filtering the data. All sequences were trimmed from right to where the average quality of 5 bp window was lower than QV25, 5 bp from the left end were trimmed, sequences with length lower than 50 bp and sequences with average quality QV25 were removed.

### Genome assembly, variant calling and global dataset

Quality filtered data further assembled using reference-based mapping with CLC Genomics Workbench 12. Mapping was done using stringent parameters with length fraction to 0.99 and similarity fraction 0.9. Mapping tracks were used to call and annotate variants. Variants with minimum allele frequency 30% with minimum coverage 10 reads were considered. For comparative analysis with the global dataset of 57,043 complete viral genomes and 974 viral genome isolates from India were downloaded from GISAID flu server (https://www.gisaid.org/), as accessed on 4^th^ July, 2020.

### Phylogenetic analysis

A total of 361 SARS-CoV-2 whole genomes sequenced in our research laboratory, as described in the above sections, were analyzed for the phylogenetic distribution. The reference genome, Wuhan/Hu-1/2019 (EPI_ISL_402125) was downloaded from GISAID flu server, which was sampled on 31^st^ Dec 2019 from Wuhan, China. Additionally, three more viral genomes were included in the phylogenetic analysis Wuhan/WH01/2019 (EPI_ISL_406798, sampled on 26 Dec 2019, Male, 44 yrs.), Wuhan/WIV04/2019 (EPI_ISL_402124, sampled on 30 Dec 2019, Female, 49 yrs.), and Wuhan/WH04/2020 (EPI_ISL_406801, sampled on 05 Jan 2020, Male, 39 yrs.). The multiple sequence alignment was performed using MAFFT (**Katoh and Standley 2013**) implemented via a phylodynamic alignment pipeline provided by Augur (https://github.com/nextstrain/augur). The subsequent alignment output files were checked, visualized and verified using PhyloSuite **(Zhang et al. 2020)**. Afterwards, the maximum likelihood phylogenetic tree was built using the Augur tree implementation pipeline with the IQ-TREE 2 **(Minh et al. 2020)** with default parameters. The selected metadata information is plotted in the time resolved phylogenetic tree was constructed using TreeTime (**Sagulenko et al. 2018**), annotated and visualized in the FigTree (**Rambaut et al., 2018**).

### Statistical analysis

The chi-square test of significance was used to check the effect of age, gender and mutations on mortality.

## Data access

The raw data generated in this study have been submitted to the NCBI BioProject database (https://www.ncbi.nlm.nih.gov/bioproject) under accession number PRJNA625669. Supplementary dataset to this manuscript are also available at Mendeley Data with DOI: 10.17632/pc38m6mwxt.1 (https://data.mendeley.com/datasets/pc38m6mwxt/draft?a=1aa66c2a-5b93-456f-816c-3f26a482dc2a). All datasets of COVID-19 are also provided on GBRC-COVID portal (http://covid.gbrc.org.in/).

## Acknowledgements

The authors are grateful to the Secretory, Department of Science and Technology (DST), and Health Commissioner Government of Gujarat, Gandhinagar, Gujarat, India. Authors are also thankful to the clinical staff for extending support in sample collection. Authors would like to acknowledge Dr. Manish Kumar, IIT-Gandhinagar for critically reviewing and providing essential inputs in the manuscript, Dr. Raghawendra Kumar and Mr. Zuber Saiyed for providing additional support to genome assembly of viral genomes.

## Author contributions

MJ, SB and CJ conceptualized the work plan and guided it for analysis of primary data, interpretation of data, and editing of the manuscript. AP, DK, AA, and MJ retrieved and analyzed the data and generated tables and figures under supervision of CJ.MJ, DK and AP wrote the manuscript. MP, JR, ZP, PT and MG did sample processing and RNA isolation. LP, KP and NS did genome sequencing. SK did data analysis and manuscript editing.

