## Supplemental_Table_S1 for "Genomic variations in SARS-CoV-2 genomes from Gujarat: Underlying role of variants in disease epidemiology"

This dataset represents additional information manuscript entitled “Genomic variations in SARS-CoV-2 genomes from Gujarat: Underlying role of variants in disease epidemiology”
**Supplementary Table S1**: The details of the samples collected from each location along with districts from the Gujarat State

| Location | District | No of Genomes |
|---|---|---|
| Ahmedabad | Ahmedabad | 125 |
| Bayad | Aravalli | 1 |
| Dhansura | Aravalli | 1 |
| Modasa | Aravalli | 14 |
| Dhanera | Banaskantha | 1 |
| Palanpur | Banaskantha | 6 |
| Bharuch | Bharuch | 1 |
| Botad | Botad | 1 |
| Dahod | Dahod | 5 |
| Dahegam | Gandhinagar | 4 |
| Gandhinagar | Gandhinagar | 19 |
| Kalol | Gandhinagar | 3 |
| Mansa | Gandhinagar | 2 |
| Kodinar | Gir Somnath | 1 |
| Una | Gir Somnath | 3 |
| Jamnagar | Jamnagar | 6 |
| Junagadh | Junagadh | 2 |
| Kapadvanj | Kheda | 1 |
| Kheda | Kheda | 2 |
| Mahemdavad | Kheda | 2 |
| Nadiad | Kheda | 2 |
| Bhuj | Kutch | 1 |
| Mandvi | Kutch | 1 |
| Kadi | Mahesana | 2 |
| Gondal | Rajkot | 1 |
| Jetpur | Rajkot | 1 |
| Rajkot | Rajkot | 16 |
| Himatnagar | Sabarkantha | 8 |
| Khedbrahma | Sabarkantha | 3 |
| Prantij | Sabarkantha | 6 |
| Talod | Sabarkantha | 1 |
| Bardoli | Surat | 2 |
| Choryasi | Surat | 9 |
| Olpad | Surat | 1 |
| Surat | Surat | 53 |
| Chuda | Surendranagar | 1 |
| Savli | Vadodara | 4 |
| Vadodara | Vadodara | 49 |
| | **Grand Total** | **361** |
