## Supplemental_Table_S3 for "Genomic variations in SARS-CoV-2 genomes from Gujarat: Underlying role of variants in disease epidemiology"

This dataset represents additional information manuscript entitled “Genomic variations in SARS-CoV-2 genomes from Gujarat: Underlying role of variants in disease epidemiology”

**Supplementary Table S3**: Lineage and Clade distribution of SARS-CoV-2 from the Districts of the Gujarat State.

| District | Lineage | | | | | | Clades | | | | | |
|---|---|---|---|---|---|---|---|---|---|---|---|---|
| | A | B | B.1 | B.1.1 | B.1.36 | B.6 | G | GH | GR | L | O | S |
| Ahmedabad | | | 51 | | 72 | 2 | 47 | 76 | | 1 | 1 | |
| Aravalli | | | 1 | | 15 | | 1 | 15 | | | | |
| Banaskantha | | | 1 | 1 | 5 | | 3 | 4 | | | | |
| Bharuch | | | 1 | | | | 1 | | | | | |
| Botad | | | | | | 1 | | | | | 1 | |
| Dahod | | | 1 | | | 4 | 1 | | | | 4 | |
| Gandhinagar | | | 14 | | 14 | | 14 | 14 | | | | |
| Gir Somnath | | | | 2 | 1 | 1 | | 1 | 2 | | 1 | |
| Jamnagar | | | 1 | | 4 | 1 | 1 | 4 | | | 1 | |
| Junagadh | | | 2 | | | | 2 | | | | | |
| Kheda | 1 | | | | 6 | | | 6 | | | | 1 |
| Kutch | | | 1 | 1 | | | 1 | | 1 | | | |
| Mahesana | | | 2 | | | | 2 | | | | | |
| Rajkot | 5 | | 5 | | 8 | | 5 | 8 | | | | 5 |
| Sabarkantha | | 1 | 10 | | 7 | | 9 | 8 | | | 1 | |
| Surat | 8 | 2 | 29 | 1 | 22 | 3 | 29 | 20 | 1 | | 8 | 7 |
| Surendranagar | | | | | 1 | | | 1 | | | | |
| Vadodara | | | 24 | | 29 | | 23 | 30 | | | | |
| **Grand Total** | **14** | **3** | **143** | **5** | **184** | **12** | **139** | **187** | **4** | **1** | **17** | **13** |
